# Bidirectional changes in excitability upon loss of both CAMK2A and CAMK2B

**DOI:** 10.1101/2022.06.07.494679

**Authors:** Martijn J. Kool, Hanna E. Bodde, Ype Elgersma, Geeske M. van Woerden

**Affiliations:** Department of Neuroscience, Erasmus MC, Wytemaweg 80, 3015 CN, Rotterdam, The Netherlands; ENCORE, Erasmus MC, Wytemaweg 80, 3015 CN, Rotterdam, The Netherlands; Department of Clinical Genetics, Erasmus MC, Wytemaweg 80, 3015 CN, Rotterdam, The Netherlands

**Author notes:** The authors declare no competing financial interests.

## Abstract

The mammalian Ca2+/calmodulin-dependent protein kinase II (CAMK2) family consists of 4 different *CAMK2* genes, encoding CAMK2A, CAMK2B, CAMK2D and CAMK2G, which have high structural homology. CAMK2A and CAMK2B are abundantly expressed in the brain; they play a unique role in proper neuronal functioning, since both CAMK2A and CAMK2B knockout mice show several behavioural and cellular phenotypes. However, our recent finding that deletion of both CAMK2A and CAMK2B is lethal indicates that they show redundancy and that the full spectrum of CAMK2 function in neurons remains to be uncovered. For example, it still remains unclear which overlapping functions are present at a single cell level in neuronal transmission and excitability.

In order to get more insight into the full spectrum of CAMK2 functions in neurons, we performed whole-cell patch clamp experiments in inducible *Camk2a/Camk2b* double knockout mice, as well as the CAMK2A and CAMK2B knockout mice. We found that whereas deletion of only CAMK2A or CAMK2B did not change excitability, simultaneous deletion of CAMK2A and CAMK2B resulted in a decrease in excitability 10 days after deletion in CA1 pyramidal neurons, which reversed to increased excitability 21 days after deletion. Additionally, loss of both CAMK2A and CAMK2B resulted in a decreased frequency of both miniature excitatory and inhibitory postsynaptic currents (mEPSC and mIPSC) 21 days after deletion, but not 10 days after deletion, an effect not seen in the single mutants. Our results indicate that CAMK2 is critically important to maintain normal excitability of hippocampal CA1 pyramidal cells, as well as normal inhibitory and excitatory synaptic transmission. Together, these results lead to new insights in how CAMK2 regulates normal neuronal function and highlight the importance of having both CAMK2A and CAMK2B expressed in high levels in the brain.

## Introduction

The mammalian Ca2+/calmodulin-dependent protein kinase II (CAMK2) family consists of 4 different *CAMK2* genes, encoding CAMK2A, CAMK2B, CAMK2D and CAMK2G, which all share high homology in their structure (Tobimatsu and Fujisawa, 1989). CAMK2A and CAMK2B are both expressed in high abundance in neurons, accounting for up to 2% of total brain protein (Erondu and Kennedy, 1985). Studies using *Camk2a* or *Camk2b* mutant mice indicate that these proteins fulfil unique functions in the brain (Silva et al., 1992a; 1992b; Hojjati et al., 2007; van Woerden et al., 2009; Borgesius et al., 2011; Kool et al., 2016). However, these isoforms may also have overlapping functions. In support for this hypothesis we recently found that simultaneous deletion of both CAMK2A and CAMK2B is lethal, indicating that critical functions of CAMK2 are masked in the single mutants due to isoform compensation and or redundancy (Kool et al., 2019). Here we set out to identify these functions at the single cell and network level.

The role of CAMK2A and CAMK2B in neuronal function and synaptic plasticity has been extensively studied using *Camk2a*^−/−^ and/or *Camk2b*^−/−^ mice (Silva et al., 1992b; van Woerden et al., 2009; Borgesius et al., 2011; Achterberg et al., 2014), as well as by studying mutant mice with knock-in missense mutations (Mayford et al., 1996; Giese et al., 1998; Elgersma et al., 2002). In addition, pharmacological approaches have been used that interfere with kinase function of both isoforms. Inhibiting CAMK2 activity in vestibular nucleus neurons (Nelson et al., 2005) and medium spiny neurons in the striatum (Klug et al., 2012) resulted in increased intrinsic excitability. Consistent with this finding, expression of constitutive active CAMK2 in CA1 pyramidal neurons decreases neuronal excitability (Varga et al., 2004), whereas the CAMK2A-T286A mouse mutant, in which the autonomous activity of CAMK2A is blocked, shows increased excitability (Sametsky et al., 2009), suggesting that CAMK2 activity is needed to regulate neuronal excitability. Indeed, blocking CAMK2 activity in cultured cortical neurons for >4 hours increases excitability and eventually leads to cell death (Ashpole et al., 2012).

At the synaptic level most studies have focussed either on CAMK2A alone or used a broad pharmacological approach to investigate the role of CAMK2 in basal synaptic transmission. For example, in hippocampal CA1 pyramidal cells and piriform cortex pyramidal cells, CAMK2 has been shown to increase the amplitude of spontaneous or miniature inhibitory postsynaptic currents (sIPSC or mIPSC), most likely through phosphorylation of the GABA_A_ receptor (Wei et al., 2004; Ghosh et al., 2015). CAMK2 mediates a glutamate-induced increase in miniature excitatory postsynaptic currents (mEPSC) frequency in cultured hippocampal neurons (Ninan and Arancio, 2004). Infusion of activated CAMK2 in hippocampal CA1 pyramidal cells also increases the amplitude of evoked EPSCs (Lledo et al., 1995), whereas knockdown of CAMK2A in dissociated hippocampal cultures decreased mEPSC amplitude, but not frequency (Barcomb et al., 2014). Only one study investigated the differential effect of CAMK2A and CAMK2B on mEPSC frequency; upon overexpression in neurons opposing effects on basal synaptic strength were found, indicating that the ratio of CAMK2A and CAMK2B is important for activity-dependent synaptic homeostasis (Thiagarajan et al., 2002). Recently another study found that CAMK2A activity is essential for maintaining basal synaptic transmission and that loss of CAMK2A, but not CAMK2B, diminishes AMPA-receptor and NMDA-receptor currents (Incontro et al., 2018).

Since the majority of the above-mentioned studies used CAMK2 inhibitors, it is difficult to disentangle whether CAMK2A, CAMK2B or both are responsible for the observed effects. In this study we set out to investigate the contribution of CAMK2A and CAMK2B, alone or in combination, on neuronal excitability and basal synaptic transmission at the single cell level. To that end, we made use of single *Camk2a^−/−^* and *Camk2b^−/−^* mutants to study the role of each of these isoforms separately, as well as inducible *Camk2a^f/f^*;*Camk2b^f/f^* knockout mice to understand how simultaneous loss of both isoforms affects neuronal function. Using whole-cell electrophysiology we found that only deletion of both CAMK2A and CAMK2B affects intrinsic excitability and basal synaptic transmission. We further show that induced deletion of both isoforms results in remarkable temporary changes. Whereas deletion of both CAMK2 isoforms initially results in decreased neuronal intrinsic excitability, this shifts to a marked increase in intrinsic excitability in just 10 days. The increase in intrinsic excitability coincides with a decrease of both miniature excitatory as well as inhibitory postsynaptic currents in the inducible *Camk2a^f/f^*;*Camk2b^f/f^* knockout mice, suggesting that the increase in excitability could be a homeostatic response of the cell to maintain its basal functions.

## Material and methods

### Animals

*Camk2a^−/−^*, *Camk2b^−/−^* and *Camk2a^f/f^*;*Camk2b^f/f^*;Cag-Cre^ER^ mice were tested in a C57Bl/6JOlaHsd background and backcrossed >16 times. Mice were genotyped between 7-10 days of age and re-genotyped after the mice were sacrificed. Genotyping records were obtained and kept by a technician not involved in the design, execution or analysis of the experiments.

Electrophysiological recordings in *Camk2a^−/−^* and *Camk2b^−/−^* mice were performed between 17 and 30 days of age. *Camk2a^f/f^*;*Camk2b^f/f^;Cag-Cre^ER^* mice were older when tested. Tamoxifen injections began at the age of 21 days until a maximum age of 30 days. Three groups were used: D10 (10 days after the first tamoxifen injection), D15 (15 days after the 1^st^ Tamoxifen injection) and D21 (21 days after the 1^st^ Tamoxifen injection). In absolute age, these time points corresponded with 31-40 days of age for mice of the D10 group, 36-45 days of age for mice of the D15 group and 42-50 days of age for mice of the D21 group.

All mice were kept group-housed in IVC cages (Sealsafe 1145T, Tecniplast) with bedding material (Lignocel BK 8/15 from Rettenmayer) on a 12/12 h light/dark cycle in 21°C (±1°C), humidity at 40-70% and with food pellets (801727CRM(P), Special Dietary Service) and water available *ad libitum*. For all experiments, mutants were compared to either WT or *Cre*–negative homozygous floxed littermates. Groups were matched for age and sex. All experiments were done during daytime and experimenters were blind for genotype throughout experiments and data analysis. All research has been performed in accordance with and approved by a Dutch Animal Ethical Committee (DEC) for animal research.

### Generation of Camk2a^−/−^, Camk2b^−/−^ and Camk2a^f/f^;Camk2b^f/f^;Cag-Cre^ER^ mice

The generation of *Camk2a^−/−^* and *Camk2b^−/−^* mice has been described previously (Elgersma et al., 2002; Kool et al., 2016) *Camk2a^f/f^*;*Camk2b^f/f^*;Cag-Cre^ER^ mice were generated by crossing *Camk2a^f/f^* (Achterberg et al., 2014) and *Camk2b^f/f^* (Kool et al., 2016) mice.

### Tamoxifen injections

*Camk2a^f/f^*;*Camk2b^f/f^* and *Camk2a^f/f^*;*Camk2b^f/f^;Cag-Cre^ER^* mice (as early as P21) were injected with 0.1mg/gr of bodyweight Tamoxifen (Sigma-Aldrich) intraperitoneally for 4 consecutive days. To keep Tamoxifen dose constant throughout injection days we kept a tight injection scheme, injecting mice 24±1 hour after the previous injection. Injections were given each day around noon. Tamoxifen was dissolved in sunflower oil (20mg/ml).

Even though Tamoxifen is not known to have an effect on physiology, we injected both *Camk2a^f/f^*;*Camk2b^f/f^* and *Camk2a^f/f^*;*Camk2b^f/f^;Cag-Cre^ER^* mice with Tamoxifen to control for any possible effects.

### Electrophysiology

Brain slices were prepared from mice using standard techniques. Briefly, mice were decapitated under deep isoflurane anaesthesia, brains were quickly removed and 320 μm thick transverse slices were cut using a vibratome (HM650V; Microm) in ice-cold cutting solution, containing (in mM): 126 Choline Chloride, 2.5 KCl, 1.25 NaH_2_PO_4_, 10 MgSO_4_, 0.5 CaCl_2_, 16.7 glucose, 26 NaHCO_3_ bubbled with 95% O_2_ and 5% CO_2_. After cutting, slices were stored in the same cutting solution for 30 seconds on 35°C and then transferred to 35°C artificial cerebrospinal fluid (ACSF) containing (in mM): 126 NaCl, 2.5 KCl, 1.25 NaH_2_PO_4_, 26 NaHCO_3_, 10 glucose, 2 MgSO_4_, 2 CaCl_2_ bubbled with 95% O_2_ and 5% CO_2_ (osmolarity between 300-310 mOsm/kg H_2_O). Afterwards, slices were kept in ACSF at room temperature for a minimum of 1 hour before onset of experiments.

Whole-cell patch-clamp recordings were made from pyramidal neurons from area CA1 in the hippocampus. Neurons were visualized using differential interference contrast (DIC) and infrared video microscopy optics on an Olympus BX51W1 microscope. Pipettes were pulled from borosilicate glass capillaries and had a resistance of 3-5 MΩ when filled with intracellular solutions containing (in mM): 120 K-gluconate, 10 KCl, 10 HEPES, 4 Mg-ATP, 10Na_2_-Phosphocreatine, 0.3 GTP (pH adjusted to 7.3 with KOH) and 0.5% biocytin (osmolarity between 275-280 mOsm/kg H_2_O). Pyramidal neurons were identified based on their firing pattern, morphology under the microscope as well as their morphology after biocytin staining. Recordings were made with a patch-clamp amplifier (Multiclamp 700B; Axon Instruments, Foster City, CA, USA). Signals were low-pass filtered at 2 kHz and digitized at 10 kHz. Only cells with series resistance under 30 MOhm were included. Series resistance was monitored during the experiments and not compensated for. Cells were rejected when series resistance changed by more than 20%. Resting membrane potential was measured in bridge mode (I=0) along with the other passive membrane properties immediately after obtaining whole-cell access. Firing pattern, action potential characteristics and threshold were analysed using Clampfit 10.3 (Axon Instruments). Physiological responses were evoked using a series of depolarizing constant-current pulses of 750 ms duration at 30 pA intervals from a holding potential of -68 mV or by using 1 ms, 1 or 1.5 nA constant-current pulses given at 10 Hz for 750 ms. Excitability was measured as the amount of action potentials elicited during the 750ms long constant-current pulse for each current step, calculated in Hertz and action potentials were identified and counted by the researcher in Clampfit. Action potential characteristics were measured on the 1^st^ and 3^rd^ action potential at the first step that induced action potentials. The action potential amplitude was calculated as the difference between threshold and peak. For rise and decay kinetics of the action potentials, maximum rise and decay slope were calculated. Action potential half width was calculated as the width of the action potential at half of the maximum amplitude. Action potential threshold was calculated taking the first derivative of the action potential, and measuring the corresponding voltage when the derivative exceeded 2 mV/ms. Membrane potentials were corrected for a (calculated) liquid junction potential of -8 mV. KN62 (Tocris Bioscience) was used in a concentration of 20 µM and autocamtide-2 related inhibitory peptide II (AIP-II; Merck Millipore) at 4 µM.

Miniature synaptic currents (mEPSCs and mIPSCs) were analysed using Mini Analysis (Synaptosoft Inc, Decatur, GA, USA). Events were detected with a threshold criterion of 3 times root-mean-square (RMS) of baseline noise. After detection, recordings were manually checked for false positive or missed events while the experimenter was still blind for genotype. mEPSCs were pharmacologically isolated using tetrodotoxin (TTX) (1 μM) and picrotoxin (50 μM). mIPSCs were isolated using TTX (1 μM), 6-cyano-7- nitroquinoxaline-2,3-dione (CNQX) (10 μM) and D,L-2-amino-5-phosphopentanoic acid (AP5; 50 μM). Cells were held at -70 mV when measuring miniature events. For mIPSCs a high chloride internal solution was used, containing (in mM): 2 NaCl, 141 KCl, 1 CaCl_2_, 10 EGTA, 2 Mg-ATP, 0.3 Na-GTP, 10 HEPES, 10 Na-phosphocreatine (pH 7.25, 295 mOsm/kg H_2_O).

### Western blot

Western blot analysis was done as described previously (Kool et al., 2016). In short, western blots were probed with primary antibodies against CAMK2A (6G9, 1:40.000, Abcam), CAMK2B (CB-β1, 1:10.000, Invitrogen) and actin (MAB1501R, 1:20.000, Chemicon) and secondary antibodies (goat anti-mouse and/or goat anti-rabbit, both 1:3000, Affinipure #115-007-003 and #111-007-003). Blots were stained and quantified using LI-COR Odyssey Scanner and Odyssey 3.0 software.

### Immunofluorescence

Immunofluorescence has been done as described previously (Kool et al., 2016).

### Data analysis and statistics

All excitability tests were analysed using a 2-way repeated measures ANOVA to determine a main effect of genotype. For all miniatures, averages were compared using a Student’s t-test. α was set at 0.05. All values represent average ± SEM. Number of mice and number of slices are depicted in the figure legends. All statistics were performed in Graphpad Prism. *p < 0.05, **p < 0.01, ***p < 0.001.

## Results

### Absence of either CAMK2A or CAMK2B does not change the intrinsic excitability of CA1 pyramidal neurons

In order to decipher the role of CAMK2A and/or CAMK2B in neuronal excitability, we measured the physiological responses upon depolarizing current stimulations in CA1 pyramidal neurons in the hippocampus of both *Camk2a^−/−^* and *Camk2b^−/−^* mice. For both mutants we found no difference in passive membrane properties such as input resistance, capacitance and resting membrane potential compared to their littermate controls (see Table 1). Additionally, we found no changes in excitability in both mutants (*Camk2a^−/−^*: effect of genotype: *F*_(1,42)_=0.06, p=0.81; *Camk2b^−/−^*: effect of genotype: *F*_(1,60)_=0.22, p=0.64; 2way repeated measures ANOVA; Figure 1).

**Figure 1:**
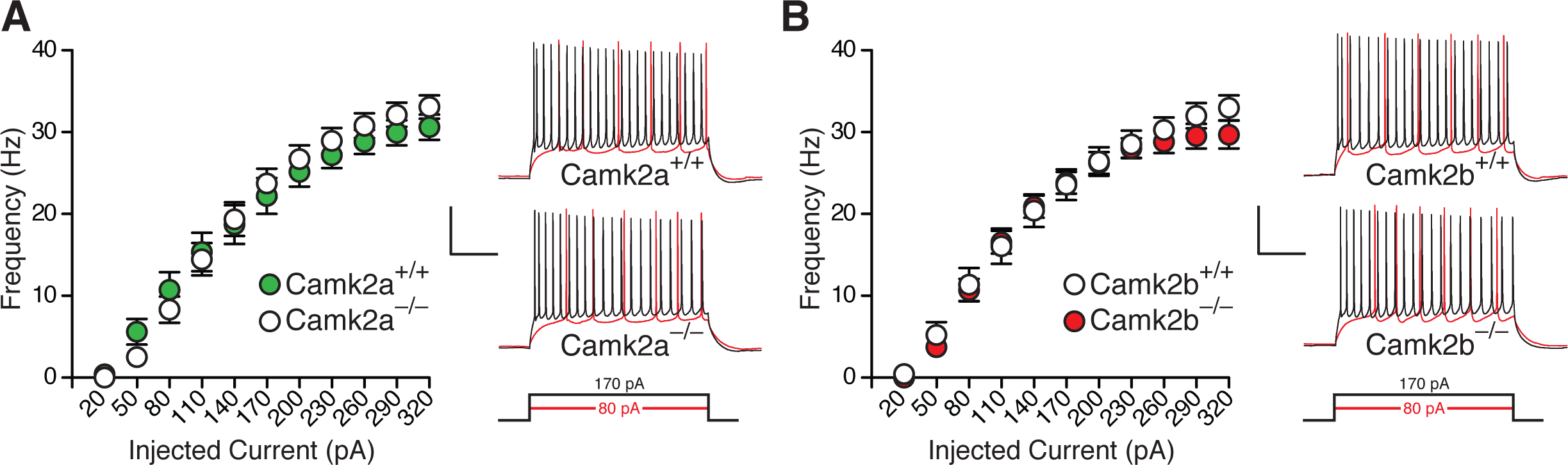
Loss of only CAMK2A and CAMK2B does not change excitability in hippocampal pyramidal CA1 neurons. **A,** Excitability is unchanged in pyramidal neurons in *Camk2a^−/−^* (n=23) compared to *Camk2a^+/+^* (n=21) mice. **B,** Excitability is unchanged in pyramidal neurons in *Camk2b^−/−^* (n=34) compared to *Camk2b^+/+^* (n=28) mice. N indicates the number of neurons measured. Scale bars indicate time (x) = 200ms and voltage (y) = 40mV.

**Table 1:** An overview of all passive membrane properties of all mice used in Figure 1 and 2. Cre+ = Camk2a^f/f^;Camk2b^f/f^;Cag-Cre^ER^. Cre- = *Camk2a*^*f/f*^;*Camk2b*^*f/f*^.

|  | *Camk2a*<sup>-/-</sup> |  |  | *Camk2b*<sup>-/-</sup> |  |  |
| --- | --- | --- | --- | --- | --- | --- |
| Parameter | WT (12) | KO (10) | p-value | WT (11) | KO (20) | p-value |
| Rm (MΩ) | 229.3±23.6 | 238.3±46.9 | ns | 220.4±30.4 | 244.4±15.6 | ns |
| Ra (MΩ) | 21.1±1.4 | 22.1±2.8 | ns | 14.4±1.4 | 17.1±0.9 | ns |
| Cm (pF) | 115.2±13.6 | 108±17.3 | ns | 110.6±10.2 | 99.2±8.8 | ns |
| Vm (mV) | -72.3±0.9 | -72.4±1.4 | ns | -71.5±1.2 | -70.4±0.9 | ns |
| tau (ms) | 2.7±0.3 | 3.3±0.7 | ns | 1.8±0.3 | 1.8±0.2 | ns |

|  | *Camk2a*<sup>f/f</sup>;*Camk2b*<sup>f/f</sup>;*CAG-Cre*<sup>ER</sup> |  |  |  |  |  |  |  |  |
| --- | --- | --- | --- | --- | --- | --- | --- | --- | --- |
|  | Day 10 |  |  | Day 15 |  |  | Day 21 |  |  |
| Parameter | Cre- (13) | Cre+ (10) | p-value | Cre- (8) | Cre+ (10) | p-value | Cre- (15) | Cre+ (18) | p-value |
| Rm (MΩ) | 240.7±9.8 | 220.6±19.3 | ns | 238.3±20.6 | 210.4±17.1 | ns | 227.6±13.2 | 252.7±12.6 | ns |
| Ra (MΩ) | 23.8±1.9 | 22.8±1.8 | ns | 24±1.6 | 25.7±1.1 | ns | 24.6±1.3 | 23.8±1.1 | ns |
| Cm (pF) | 90.8±7.4 | 91.9±13.2 | ns | 72.6±8.2 | 85.5±3.9 | ns | 87.2±5.3 | 82.6±5.1 | ns |
| Vm (mV) | -72.7±1.8 | -71.0±1.7 | ns | -74±2.1 | -73.9±2.3 | ns | -74.1±0.7 | -76.3±1.4 | ns |
| tau (ms) | 2.4±0.3 | 2.7±0.3 | ns | 2.5±0.4 | 2.7±0.3 | ns | 2.5±0.2 | 2.2±0.2 | ns |

|  | *Camk2a*<sup>f/f</sup>;*Camk2b*<sup>f/f</sup>;*CAG-Cre*<sup>ER</sup> |  |  |  |  |  |
| --- | --- | --- | --- | --- | --- | --- |
|  | OIL D10 |  |  | OIL D21 |  |  |
| Parameter | Cre- (13) | Cre+ (14) | p-value | Cre- (12) | Cre+ (13) | p-value |
| Rm (MΩ) | 241.5±15.4 | 227.9±15.2 | ns | 239.1±15.3 | 243.3±12.3 | ns |
| Ra (MΩ) | 24.3±1.4 | 25.4±1.2 | ns | 24.9±1.3 | 21.8±1.8 | ns |
| Cm (pF) | 84.7±5.4 | 80.5±4.3 | ns | 94.1±8.1 | 75.8±4.3 | ns |
| Vm (mV) | -71.2±1.3 | -71.5±1.3 | ns | -70.5±1.4 | -72.6±1.1 | ns |
| tau (ms) | 2.4±0.2 | 2.6±0.2 | ns | 3.1±0.3 | 2.1±0.3 | 0.0164 |

### Absence of both CAMK2A and CAMK2B causes a bidirectional change in excitability over time

Since we recently showed clear redundancy in some functions for CAMK2A and CAMK2B (Kool et al., 2019), we considered the possibility that also at the level of neuronal excitability, loss of CAMK2A is compensated for by CAMK2B and *vice versa*. Therefore, we decided to measure excitability in mice lacking both CAMK2A and CAMK2B. Because *Camk2a^−/−^*;*Camk2b^−/−^* mice die at P1, we made use of our inducible homozygous floxed *Camk2a^f/f^*;*Camk2b^f/f^* mice (Kool et al., 2019). Simultaneous deletion of *Camk2a* and *Camk2b* in adult mice results in lethality 24-53 days after onset of deletion. Hence we decided to measure the neuronal changes over time in the three weeks following gene deletion but before the mice become clearly affected. We injected *Camk2a^f/f^*;*Camk2b^f/f^;Cag-Cre^ER^* mice with Tamoxifen from postnatal day 21 (P21) onwards for 4 consecutive days and measured neuronal excitability on three different time points: 10 days (D10), 15 days (D15) and 21 days (D21) after the 1^st^ injection. Western blot analysis of prefrontal cortex tissue of the brains used for the whole cell experiments showed a 60% decrease of CAMK2A and an 85% decrease of CAMK2B protein on D10, which was further decreased on D15. Maximal reduction of protein level was obtained on D21 with <14% of CAMK2A and <5% of CAMK2B left in lysates (Figure 2A). First, we compared passive membrane properties, including input resistance, capacitance and resting membrane potential, of hippocampal CA1 pyramidal neurons at the different time points. Overall, we detected no differences in passive membrane properties on D10, D15 and D21 (see Table 1). Next, we measured voltage responses in CA1 pyramidal neurons upon depolarizing current injections. On D10 *Camk2a^f/f^*;*Camk2b^f/f^;Cag-Cre^ER^* mice showed a *decrease* in excitability (effect of genotype: *F*_(1,21)_=7.64, p=0.012; Figure 2B). To our surprise, when measuring pyramidal neurons on D15, excitability was no longer significantly different between the *Camk2a^f/f^*;*Camk2b^f/f^;Cag-Cre^ER^* mutants and controls (effect of genotype: *F*_(1,16)_=0.02, p=0.89; 2way repeated measures ANOVA; Figure 2C), whereas on D21 *Camk2a^f/f^*;*Camk2b^f/f^;Cag-Cre^ER^* pyramidal neurons showed an *increase* in excitability compared to controls (effect of genotype: *F*_(1,31)_=15.25, p<0.001; 2-way repeated measures ANOVA; Figure 2D). Importantly, *Camk2a^f/f^*;*Camk2b^f/f^;Cag-Cre^ER^* mice injected with vehicle (sunflower oil without Tamoxifen) showed no changes in excitability both 10 and 21 days after the first vehicle injection (D10: effect of genotype: *F*_(1,25)_=0.94, p=0.34; D21: effect of genotype: *F*_(1,23)_=0.95, p=0.34; 2-way repeated measures ANOVA; Figure 2E and F). Taken together, whereas excitability was not changed in *Camk2a^−/−^* and *Camk2b^−/−^* single knockout mice, simultaneous deletion of both CAMK2A and CAMK2B initially induced a *decrease* in excitability followed by an *increase* in excitability.

**Figure 2:**
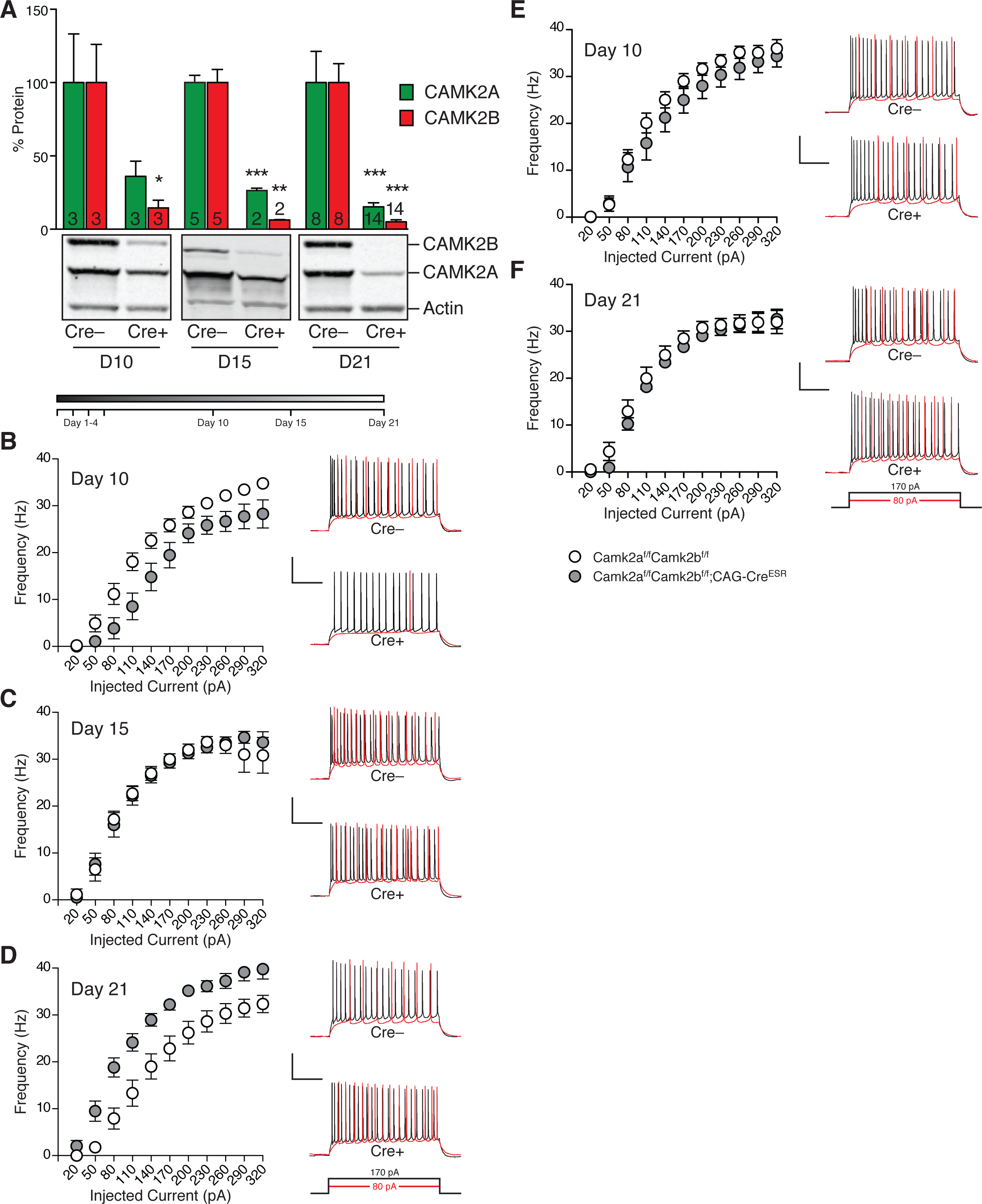
Loss of both CAMK2A and CAMK2B causes a bidirectional change in excitability in time **A,** CAMK2A and CAMK2B protein quantification using Western Blot of prefrontal cortex of *Camk2a^f/f^;Camk2b^f/f^-CAG-Cre^ER^* mice (Cre+) versus *Camk2a^f/f^;Camk2b^f/f^* mice (Cre–) 10, 15 and 21 days after the first Tamoxifen injection. Quantification of all Cre+ samples was compared to normalized Cre– samples from the same age (set at 100%). Statistics: Cre- vs Cre+: CAMK2A; D10: t=1.8, p=0.14; D15: t=9.2, p<0.001; D21: t=5.2, p<0.0001; CAMK2B; D10: t=3.2], p=0.032; D15: t=6.3, p<0.01; D21: t=9.8, p<0.0001; unpaired two tailed t-test. Actin was used as a loading control and number of samples is depicted in the bars. **B,** Cre+ mice (n=10) show decreased excitability 10 days after onset of gene deletion compared to Cre– mice (n=13). **C,** Cre+ mice (n=10) show no change in excitability 15 days after onset of gene deletion compared to Cre– mice (n=8). **D,** Cre+ mice (n=18) show increased excitability 21 days after onset of gene deletion compared to Cre– mice (n=15). **E,** Cre+ mice (n=14) show no change in excitability 10 days after injection of vehicle compared to Cre– mice (n=13). **F,** Cre+ mice (n=13) show no change in excitability 21 days after injection of vehicle compared to Cre– mice (n=12). N indicates the number of neurons measured, except for **A**. Scale bars in **B-F** indicate time (x) = 200ms and voltage (y) = 40mV.

CAMK2 is known to have both a catalytic as well as a structural role in neurons (Giese et al., 1998; Elgersma et al., 2002; Hojjati et al., 2007; Borgesius et al., 2011). To distinguish between the structural and catalytic role of CAMK2 in the observed phenotypes, excitability was measured again in CA1 pyramidal neurons of *Camk2a^f/f^*;*Camk2b^f/f^;Cag- Cre^ER^* mice; this time not after injection with Tamoxifen, but in the presence or absence of two types of CAMK2 inhibitors: the calmodulin-binding competitor KN62 and substrate-binding competitor AIP-II (autocamtide-2 related inhibitory peptide II). Neither wash-in of 20uM KN62, nor 2.5h incubation of AIP-II affected excitability in CA1 pyramidal cells (KN62: effect of genotype: F_(1,26)_=0.24, p=0.63; AIP-II: effect of genotype: *F*_(1,20)_=0.08, p=0.79; 2-way repeated measures ANOVA; Figure 3A and B), suggesting that the changes in excitability are not due to acute loss of CAMK2 activity.

**Figure 3:**
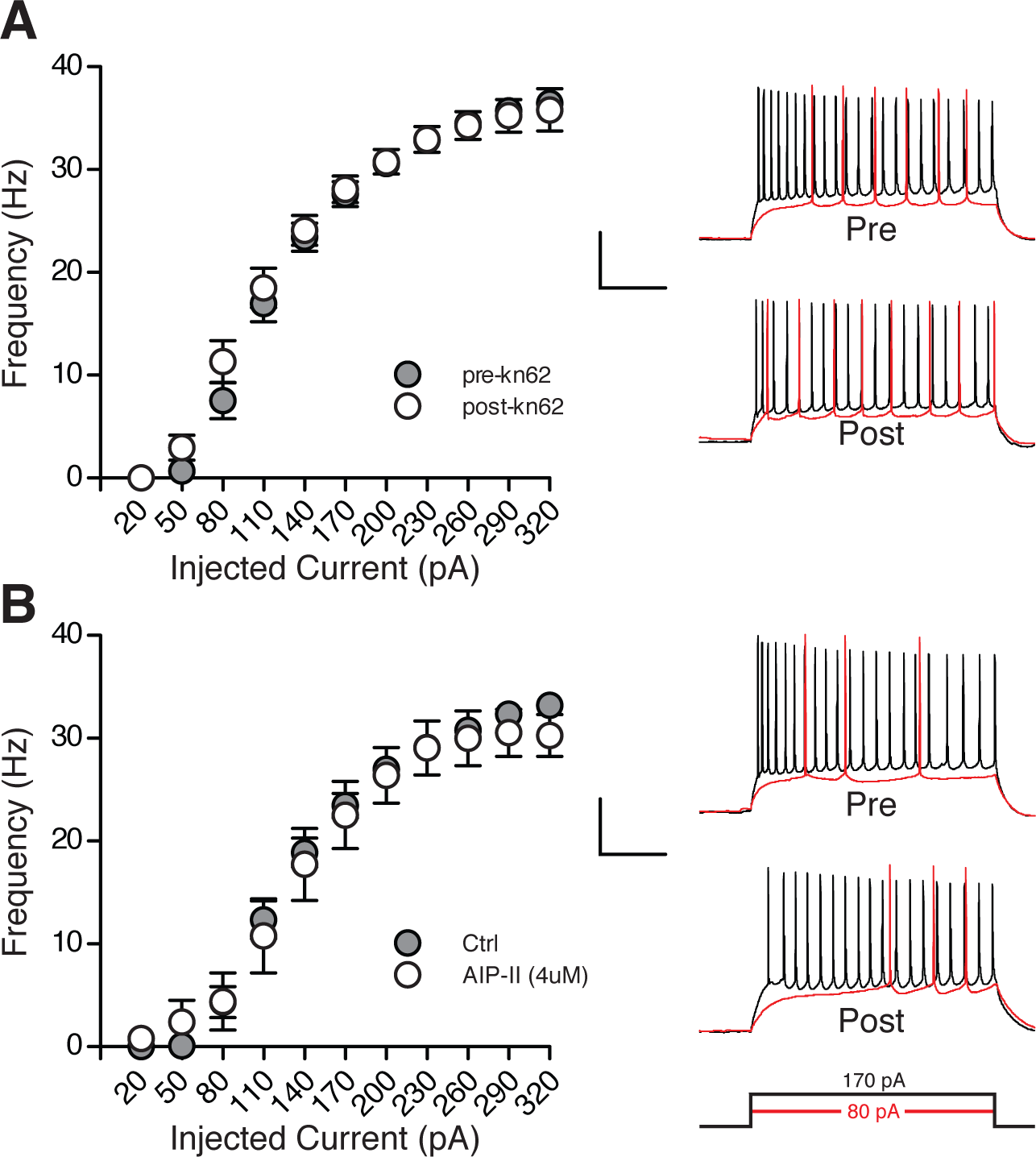
Blocking CAMK2 activity for several hours does not change excitability. **A,** KN62 wash-in for 10 minutes does not change excitability in CA1 pyramidal (n=14) compared to control conditions with no KN62 (n=14). **B,** Incubation of Autocamtide-2 related inhibitory peptide II (AIP-II) for 2.5 hours does not change excitability in CA1 pyramidal neurons (n=10) compared to control conditions with no AIP-II (n=12). In these experiments *Camk2a^f/f^*;*Camk2b^f/f^;Cag-Cre^ER^* mice who received no Tamoxifen injections were used and were therefore used as controls. N indicates the number of neurons measured. Scale bars indicate time (x) = 200ms and voltage (y) = 40mV.

### Absence of CAMK2 changes action potential threshold

Changes in excitability could result from changes in the active properties of CA1 neurons. Thus we analysed the action potential properties including the threshold, maximum amplitude, rise and decay slopes, and half-width. We analysed the properties of the first and the third action potential at the first step that induced action potentials from mice on D10 and D21. We found no changes in amplitude, maximum rise slope, maximum decay slope or action potential half-width on D10 (Figure 4). However, the threshold at which the action potential was generated on D10 for the first and third action potential was significantly more positive (Figure 4D). However, 21 days after onset of deletion, we found that mice lacking CAMK2 had a significantly more negative threshold, with no changes in amplitude, maximum rise slope, maximum decay slope or action potential half-width (Figure 5). Also, using a more physiological approach to elicit action potentials using high current squared pulses for 1 ms at 10Hz, we still found a more negative threshold in *Camk2a^f/f^*;*Camk2b^f/f^;Cag-Cre^ER^* mice on D21 (Cre- vs Cre+: t=2.3, p=0.29; unpaired two tailed t-test; Figure 5G). Taken together, loss of CAMK2 significantly changed the action potential threshold first to a more positive level, which then shifted to a more negative level.

**Figure 4:**
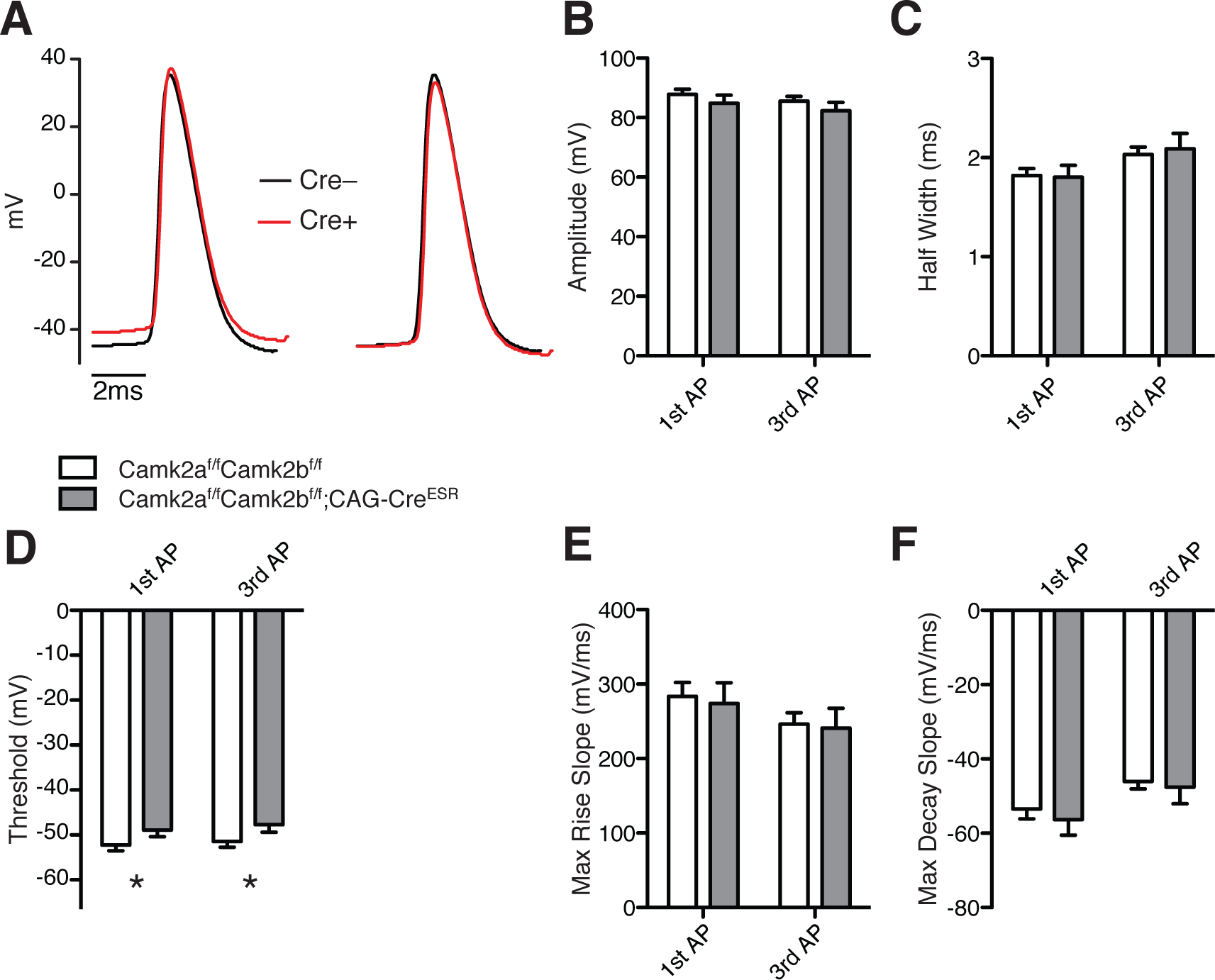
Action potential characteristics 10 days after onset of gene deletion of *Camk2a* and *Camk2b* in *Camk2a^f/f^*;*Camk2b^f/f^;Cag-Cre^ER^* mice. **A,** (*Left*) Average action potential for *Camk2a^f/f^*;*Camk2b^f/f^;Cag-Cre^ER^* mice (Cre+; red) and *Camk2a^f/f^;Camk2b^f/f^* mice (Cre–; black). Averages were taken from the first action potential at the first depolarizing step that induced action potentials. Scale bar indicates time (x) = 2ms and absolute voltage is shown on the y-axis. (*Right*) Same averages as left, but now overlapped to allow easier comparison for overall shape. **B,** No difference in amplitude of action potentials in the first and third action potential between Cre+ and Cre– mice. **C,** No difference in the half width of action potentials in the first and third action potential between Cre+ and Cre– mice. **D,** Cre+ mice show a significantly more depolarized action potential threshold in the first and third action potential compared to Cre– mice. **E,** No difference in maximum rise slope of action potentials in the first and third action potential between Cre+ and Cre– mice. **F,** No difference in maximum decay slope of action potentials in the first and third action potential between Cre+ and Cre– mice. For all graphs: Cre+ (n=10), Cre– (n=13). N indicates number of neurons measured.

**Figure 5:**
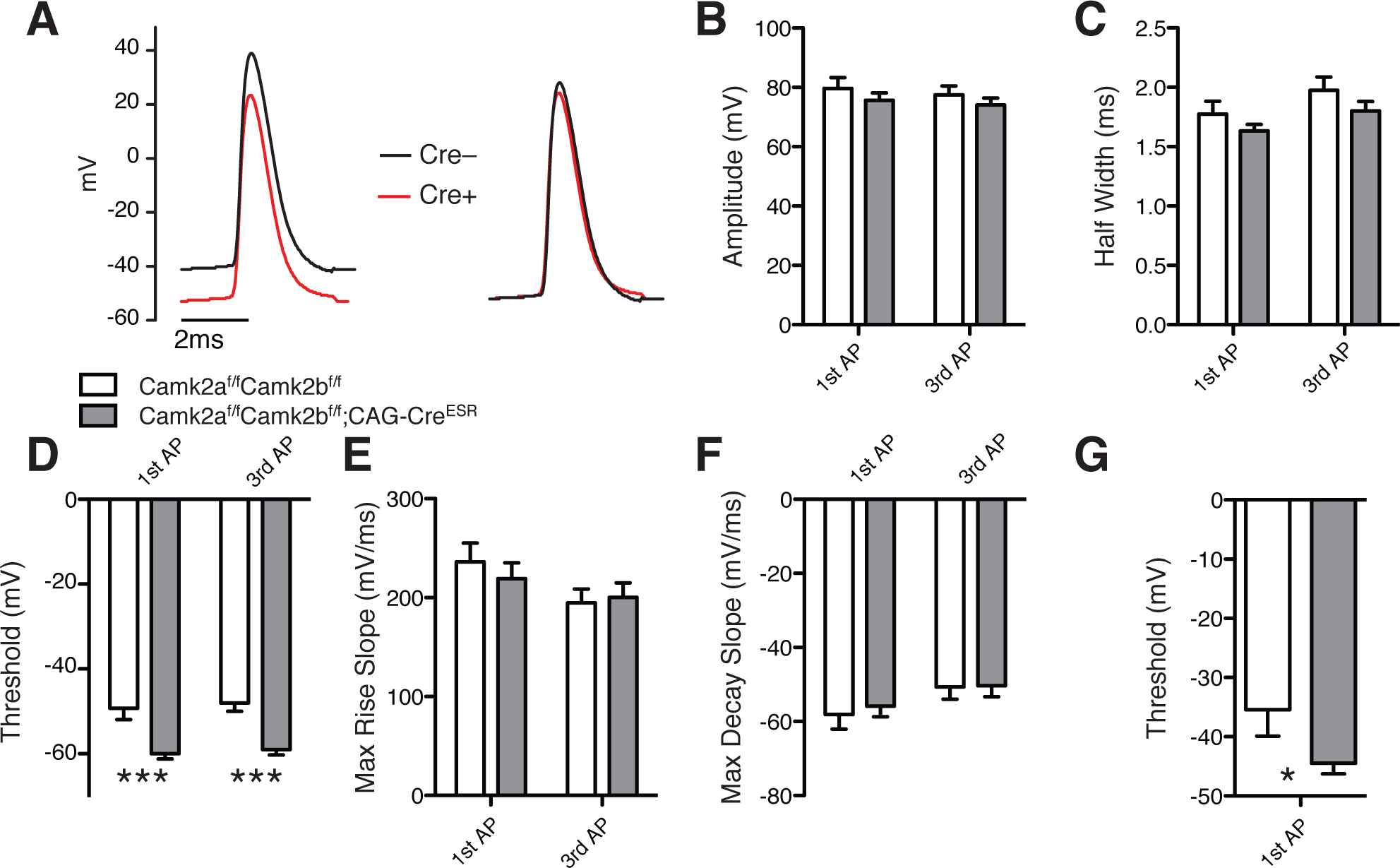
Action potential characteristics 21 days after onset of gene deletion of *Camk2a* and *Camk2b*. **A,** (*Left*) Average action potential for *Camk2a^f/f^*;*Camk2b^f/f^;Cag-Cre^ER^* mice (Cre+; red) and *Camk2a^f/f^;Camk2b^f/f^* mice (Cre–; black). Averages were taken from the first action potential from the first depolarizing step that induced action potentials. Scale bar indicates time (x) = 2ms and absolute voltage is shown on the y-axis. (*Right*) Same averages as left, but now overlapped to allow easier comparison for overall shape. **B,** No difference in amplitude of action potentials in the first and third action potential between Cre+ and Cre– mice. **C,** No difference in the half width of action potentials in the first and third action potential between Cre+ and Cre– mice. **D,** Cre+ mice show a significantly more hyperpolarized action potential threshold in the first and third action potential compared to Cre– mice. **E,** No difference in maximum rate of rise of the first or the third action potential between Cre+ and Cre– mice. **F,** No difference in maximum decay slope of action potentials in the first and third action potential between Cre+ and Cre– mice. For all graphs (**A**-**F**): Cre+ (n=18), Cre– (n=11). **G,** a more physiological protocol using 1ms squared pulses at 10Hz instead of continuous depolarization still show a significantly more hyperpolarized action potential threshold in Cre+ (n=22) compared to Cre– (n=9). N indicates number of neurons measured.

### Reduction of both inhibitory and excitatory inputs upon loss of CAMK2

Hypo- and hyperexcitability of neurons is often caused by changes in the inhibition to excitation ratio (Isaacson and Scanziani, 2011; Zhou and Yu, 2018). CAMK2A and CAMK2B are highly expressed at the synapse, regulating functional as well as structural plasticity (Lee et al., 2009; Kim et al., 2015) and changes in the ratio of CAMK2A and CAMK2B are shown to influence unitary synaptic strength as well as mEPSC frequency (Thiagarajan et al., 2002). Thus, it is plausible that loss of one or both CAMK2 isoforms affects the inhibitory and/or excitatory input a neuron receives, resulting in changes in excitability. Therefore, we measured both miniature excitatory and inhibitory postsynaptic currents (mEPSCs and mIPSCs) in hippocampal pyramidal cells of both *Camk2a^−/−^* and *Camk2b^−/−^* mice. Importantly, mEPSC and mIPSC frequency and amplitude were unaffected in both *Camk2a^−/−^* and *Camk2b^−/−^* mice (*Camk2a^−/−^*: mEPSC frequency: t=0.01, p=0.99; mEPSC amplitude: t=0.27, p=0.79; mIPSC frequency: t=0.05, p=0.96; mIPSC amplitude: t=0.51, p=0.62; *Camk2b^−/−^*: mEPSC frequency: t=0.37, p=0.71; mEPSC amplitude: t=0.78, p=0.45; mIPSC frequency: t=1.98, p=0.06; mIPSC amplitude: t=0.66, p=0.51; unpaired two tailed t-test; Figure 6). We then hypothesized that redundancy could play a role on the level of mEPSC and mIPSC frequency and amplitude similar as in excitability. We measured *Camk2a^f/f^*;*Camk2b^f/f^;Cag-Cre^ER^* mice on the two time points at which excitability was changed: 10 days (D10) and 21 days (D21) after the 1^st^ injection. Surprisingly, on D10, mEPSCs and mIPSCs showed no difference in frequency and amplitude between *Camk2a^f/f^*;*Camk2b^f/f^;Cag-Cre^ER^* mice and *Camk2a^f/f^*;*Camk2b^f/f^* control littermates (Cre- vs Cre+: mEPSC frequency: t=0.65, p=0.52; mEPSC amplitude: t=1.66, p=0.11; mIPSC frequency: t=1.0, p=0.31; mIPSC amplitude: t=0.32, p=0.75; unpaired two tailed t-test; Figure 7A and B). On D21 however, we found a significant decrease in mEPSC and mIPSC frequency but not in amplitude (Cre- vs Cre+: mEPSC frequency: t=2.14, p=0.0398; mEPSC amplitude: t=0.71, p=0.48; mIPSC frequency: t=3.49, p<0.01; mIPSC amplitude: t=1.2, p=0.22; unpaired two tailed t-test; Figure 7C and D). To understand whether the reduced frequency was due to loss of synapses, neurons were filled with biocytin and the number of spines as well as their shape was analysed. Whereas there was no change in the density of spines per 10 micron length of dendrite, primary dendrites, branches and total neurite length (number of spines: t=0.06, p=0.95; unpaired two tailed t-test; Figure 8A, C-E), we did find a trend towards more immature spines (filopodia) in the *Camk2a^f/f^*;*Camk2b^f/f^;Cag-Cre^ER^* mice (filopodia vs. spines: t=3.29, p=0.046; unpaired two tailed t-test; Figure 8B). Taken together, these results imply that the hypo-excitability seen in the *Camk2a^f/f^*;*Camk2b^f/f^;Cag-Cre^ER^* mice at day 10 after induction of gene deletion, is not caused by initial changes in the inhibition to excitation ratio.

**Figure 6:**
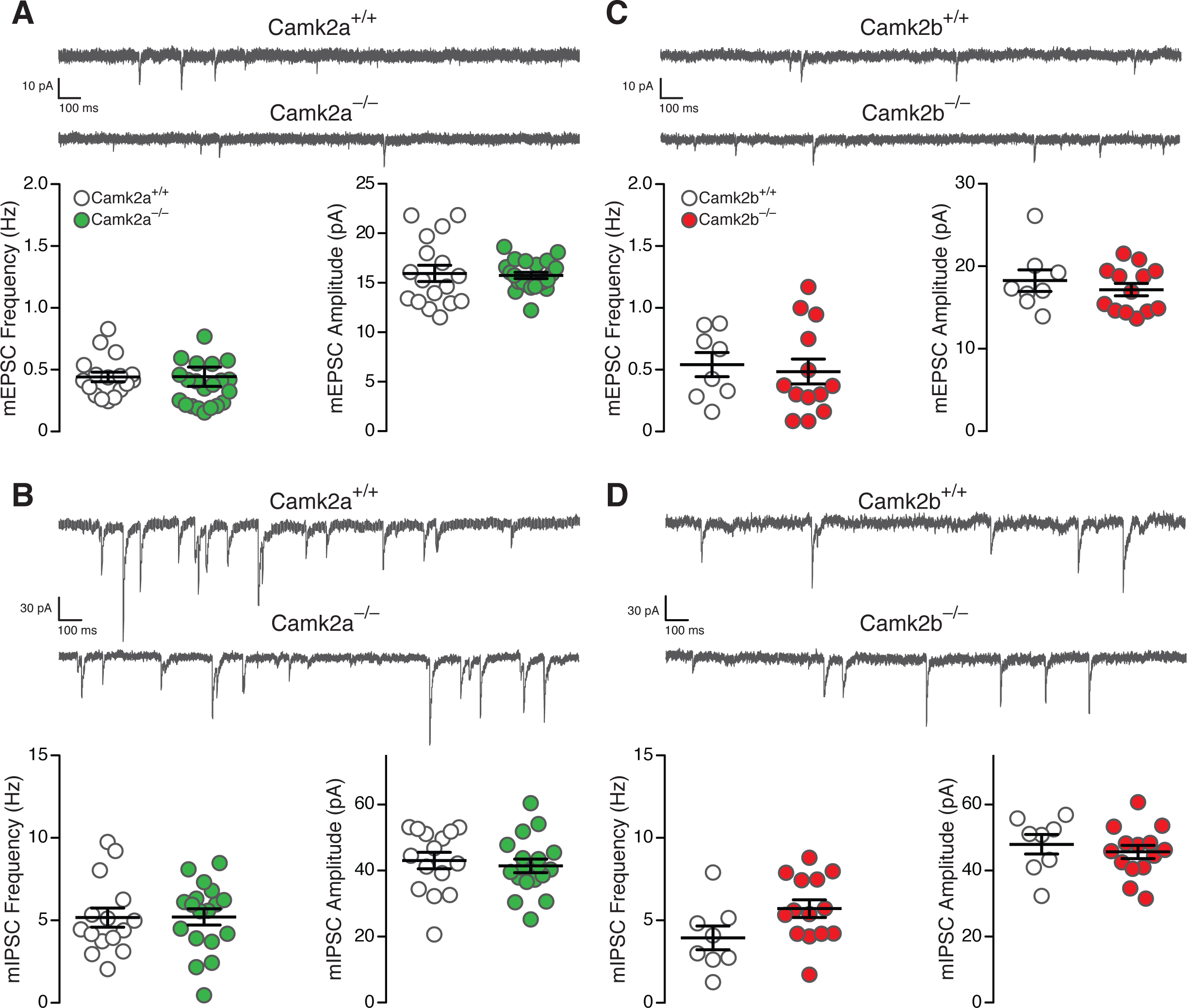
No changes in mEPSC and mIPSC frequency and amplitude of *Camk2a^−/−^* and *Camk2b^−/−^*. **A,** Representative traces (top) and frequency and amplitude (bottom) of miniature excitatory postsynaptic currents (mEPSC) measured from CA1 pyramidal cells. *Camk2a^−/−^* mice (n=21) show no change in frequency (*Camk2a^+/+^* vs *Camk2a^−/−^*: 0.44 vs 0.44 Hz) and amplitude (*Camk2a^+/+^* vs *Camk2a^−/−^*: 15.9 vs 15.7 pA) compared to *Camk2a^+/+^* mice (n=17). **B,** Representative traces (top) and frequency and amplitude (bottom) of miniature inhibitory postsynaptic currents (mIPSC) measured from CA1 pyramidal cells. *Camk2a^−/−^* mice (n=18) show no change in frequency (*Camk2a^+/+^* vs *Camk2a^−/−^*: 5.17 vs 5.21 Hz) and amplitude (*Camk2a^+/+^* vs *Camk2a^−/−^*: 43.03 vs 41.42 pA) compared to *Camk2a^+/+^* mice (n=15). **C,** Representative traces (top) and frequency and amplitude (bottom) of miniature excitatory postsynaptic currents (mEPSC) measured from CA1 pyramidal cells. *Camk2b^−/−^* mice (n=13) show no change in frequency (*Camk2b^+/+^* vs *Camk2b^−/−^*: 0.54 vs 0.50 Hz) and amplitude (*Camk2b^+/+^* vs *Camk2b^−/−^*: 18.26 vs 17.03 pA) compared to *Camk2b^+/+^* mice (n=8). **D,** Representative traces (top) and frequency and amplitude (bottom) of miniature inhibitory postsynaptic currents (mIPSC) measured from CA1 pyramidal cells. *Camk2b^−/−^* mice (n=14) show no change in frequency (*Camk2b^+/+^* vs *Camk2b^−/−^*: 3.95 vs 5.71 Hz) and amplitude (*Camk2b^+/+^* vs *Camk2b^−/−^*: 47.97 vs 45.65 pA) compared to *Camk2b^+/+^* mice (n=8). Scale for all mEPSCs indicates time (x) = 100ms and current (y) = 10pA: Scale for all mIPSCs indicates time (x) = 100ms and current (y) = 30pA. N indicates number of neurons measured.

**Figure 7:**
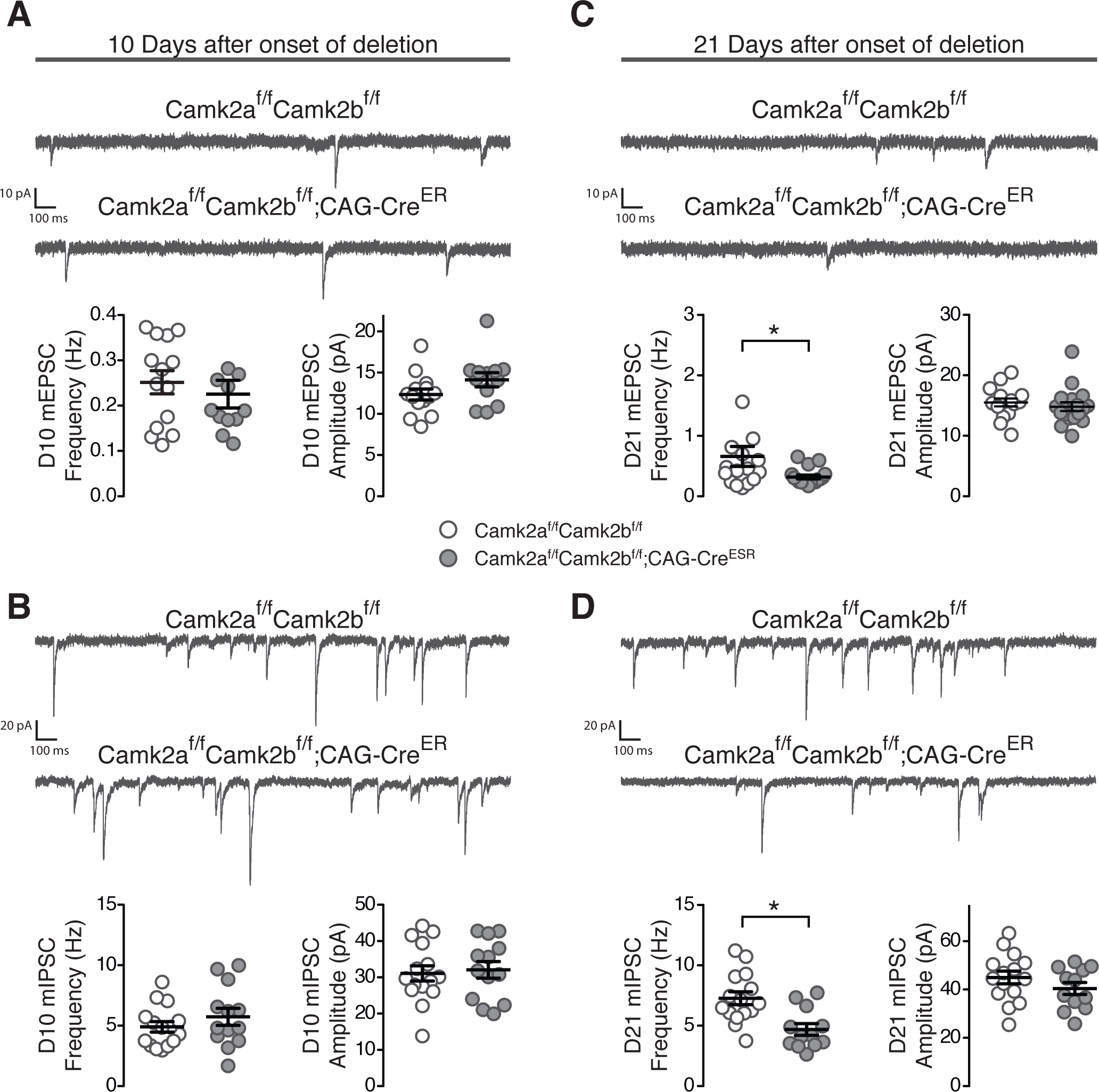
A decrease in mEPSC and mIPSC frequency 21 days after onset of gene deletion of *Camk2a* and *Camk2b*. **A,** Representative traces (top) and frequency and amplitude (bottom) of miniature excitatory postsynaptic currents (mEPSC) measured from CA1 pyramidal cells 10 days after onset of gene deletion. *Camk2a^f/f^*;*Camk2b^f/f^;Cag-Cre^ER^* mice (Cre+; n=11) show no change in frequency (Cre+ vs Cre–: 0.23 vs 0.25 Hz) and amplitude (Cre+ vs Cre–: 14.13 vs 12.35 pA) compared to *Camk2a^f/f^;Camk2b^f/f^* mice (Cre–; n=14). **B,** Representative traces (top) and frequency and amplitude (bottom) of miniature inhibitory postsynaptic currents (mIPSC) measured from CA1 pyramidal cells 10 days after onset of gene deletion. *Camk2a^f/f^*;*Camk2b^f/f^;Cag-Cre^ER^* mice (Cre+; n=13) show no change in frequency (Cre+ vs Cre–: 5.73 vs 4.90 Hz) and amplitude (Cre+ vs Cre–: 32.1 vs 31.1 pA) compared to *Camk2a^f/f^;Camk2b^f/f^* mice (Cre–; n=15). **C,** Representative traces (top) and frequency and amplitude (bottom) of mEPSCs measured from CA1 pyramidal cells 21 days after onset of gene deletion. *Camk2a^f/f^*;*Camk2b^f/f^;Cag-Cre^ER^* mice (Cre+; n=18) show a decrease in frequency (Cre+ vs Cre–: 0.32 vs 0.66 Hz) but not in amplitude (Cre+ vs Cre–: 14.82 vs 15.51 pA) of mEPSCs compared to *Camk2a^f/f^;Camk2b^f/f^* mice (Cre–; n=16). **D,** Representative traces (top) and frequency and amplitude (bottom) of mIPSCs measured from CA1 pyramidal cells 21 days after onset of gene deletion. *Camk2a^f/f^*;*Camk2b^f/f^;Cag-Cre^ER^* mice (Cre+; n=12) show a decrease in frequency (Cre+ vs Cre–: 4.70 vs 7.28 Hz) but not in amplitude (Cre+ vs Cre–: 40.40 vs 44.97 pA) of mIPSCs compared to *Camk2a^f/f^;Camk2b^f/f^* mice (Cre–; n=15). Scale for all mEPSCs indicates time (x) = 100ms and current (y) = 10pA: Scale for all mIPSCs indicates time (x) = 100ms and current (y) = 20pA. N indicates number of neurons measured.

**Figure 8:**
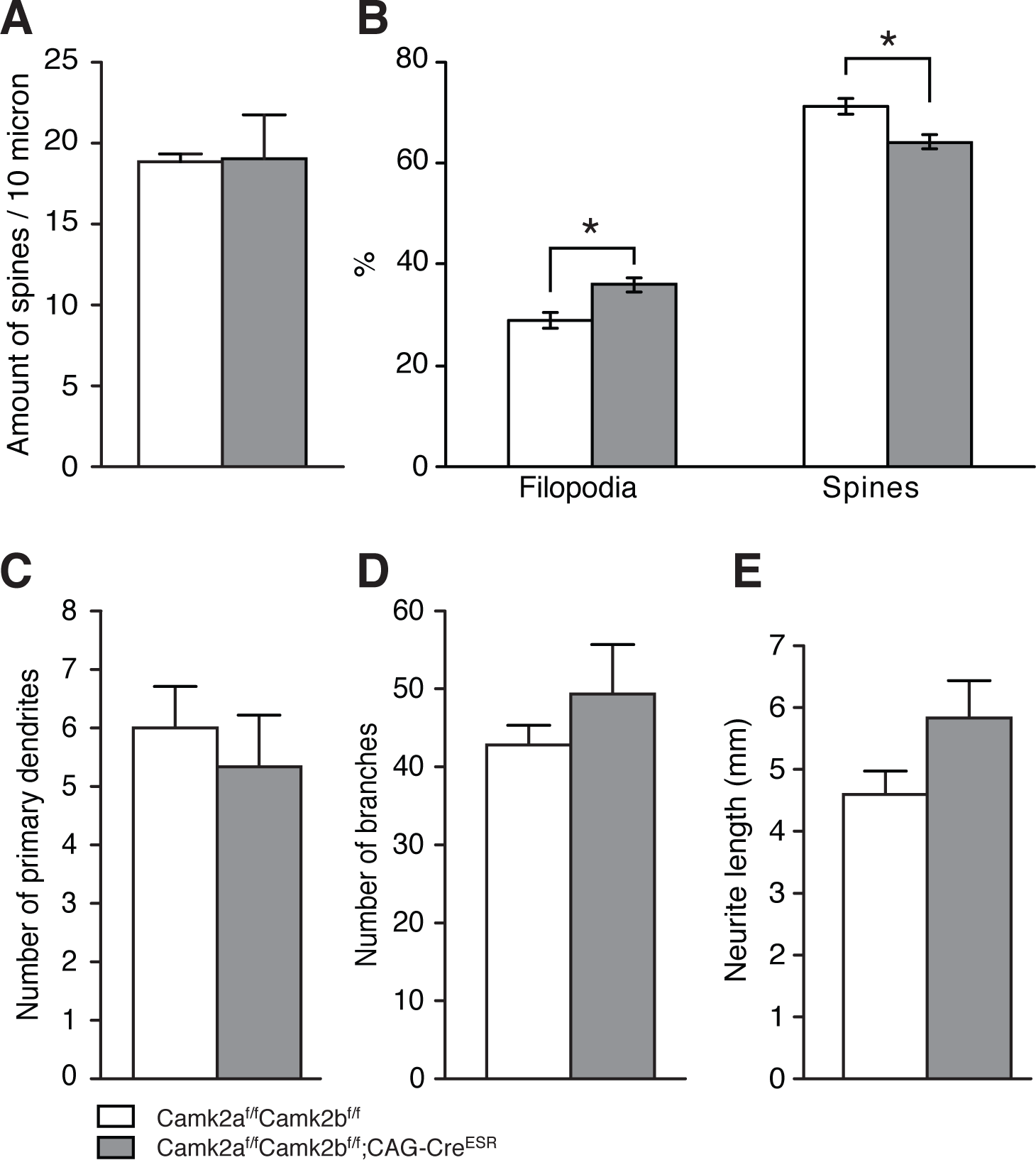
Loss of CAMK2A and CAMK2B causes an increase of immature spines (filopodia). **A,** No change in the density of spines per 10 micron length of dendrite in *Camk2a^f/f^*;*Camk2b^f/f^;Cag-Cre^ER^* mice compared to *Camk2a^f/f^*;*Camk2b^f/f^* mice. **B,** *Camk2a^f/f^*;*Camk2b^f/f^;Cag-Cre^ER^* mice show significantly more immature spines than *Camk2a^f/f^*;*Camk2b^f/f^* mice. For **A** and **B** 544 spines/filopodia from 4 neurons from 3 different *Camk2a^f/f^*;*Camk2b^f/f^;Cag-Cre^ER^* mice were used versus 540 spines/filopodia from 3 neurons from 3 different *Camk2a^f/f^*;*Camk2b^f/f^* mice. **C,** No change in the number of primary dendrites in *Camk2a^f/f^*;*Camk2b^f/f^;Cag-Cre^ER^* mice compared to *Camk2a^f/f^*;*Camk2b^f/f^* mice. **D,** No change in the number of neuronal branches in *Camk2a^f/f^*;*Camk2b^f/f^;Cag-Cre^ER^* mice compared to *Camk2a^f/f^*;*Camk2b^f/f^* mice. **E,** No change in the total neurite length in *Camk2a^f/f^*;*Camk2b^f/f^;Cag-Cre^ER^* mice compared to *Camk2a^f/f^*;*Camk2b^f/f^* mice. For graphs **C-E** 3 neurons were analysed from 3 different *Camk2a^f/f^*;*Camk2b^f/f^;Cag-Cre^ER^* mice and 5 neurons from 4 different *Camk2a^f/f^*;*Camk2b^f/f^* mice.

## Discussion

Here we made use of the *Camk2a^−/−^*, the *Camk2b^−/−^* as well as the *Camk2a^f/f^*;*Camk2b^f/f^;Cag- Cre^ER^* mice to study the effect of loss of either CAMK2A or CAMK2B or both simultaneously on neuronal excitability in hippocampal CA1 pyramidal neurons. Whereas loss of only CAMK2A or CAMK2B did not change the excitability, simultaneous loss of CAMK2A and CAMK2B resulted in a bidirectional change in excitability over time. Upon acute loss of both CAMK2 isoforms, excitability of hippocampal CA1 pyramidal cells *decreased,* but over time this was converted to an *increase* in excitability. These changes are accompanied by an initial increase of action potential threshold followed by a decrease in the threshold for which the underlying mechanisms are unknown. The changes in excitability seem not to be caused by changes in inhibition and excitation, since initially these inputs are not affected in our mini analysis; only at the latest time point a decrease was found in the frequency of both inhibitory and excitatory synaptic inputs.

It is interesting that an effect on excitability was observed only when CAMK2A and CAMK2B are both absent. This indicates that either one of them can fulfill the function needed to maintain normal intrinsic neuronal excitability. Several studies showed involvement of the kinase activity of CAMK2 in neuronal excitability, most of which indicate that CAMK2 functions to inhibit neuronal excitability. In vestibular nucleus neurons, loss of CAMK2 activity results in an increase of intrinsic excitability mediated through BK-type calcium-activated potassium channels (Nelson et al., 2005). In Drosophila, CAMK2 was found to phosphorylate the *ether à go-go* (EAG) potassium channel, and inhibition of CAMK2 results in hyper-excitability (Wang et al., 2002). The K_v_4.2 channel is also a substrate for CAMK2, and increased CAMK2 activity results in a decrease in excitability (Varga et al., 2004). In cardiac tissue and the axon initial segment of Purkinje cells, ý_IV_-spectrin-dependent targeting of CAMK2 to Na_v_1.5 allows for phosphorylation of Na_v_1.5 on S571, altering excitability (Hund et al., 2010).

Surprisingly, we find a bidirectional change in excitability over time, starting with hypo- excitability. There are several explanations for this switch: i) dose-dependence; we find at 10 days after gene deletion still 40% of CAMK2A and 15% of CAMK2B left in the prefrontal cortex. Since we have recently shown that the half-life in the hippocampus is slightly longer than in the cortex (Kool et al., 2019), the expression levels of CAMK2 in the hippocampus at this time point are likely somewhat higher than what we measured in the prefrontal cortex. This would mean that in total a fair amount of the kinase is still present when we see the decrease in excitability. However, the *Camk2a^−/−^* or *Camk2b^−/−^* mice, in which there is still 50% of CAMK2 present, did not show any change in excitability. It is possible that adult deletion in our mouse model prohibits the neuron to compensate for loss of 50% of the most abundant protein, possibly showing the direct involvement of CAMK2 in neuronal excitability. Upon further decrease of CAMK2 levels, perhaps the neuron tries to compensate for the total loss of CAMK2 and the accompanying loss of input on the synapses by homeostatically increasing the excitability at the soma. ii) subcellular localization of CAMK2 loss; CAMK2 is present in the soma, in dendrites as well as in spines. Additionally, there is a pool of CAMK2A mRNA present in the dendrites (Burgin et al., 1990; Benson et al., 1992). Depending on which pool of CAMK2 is depleted first upon acute gene deletion, different channels and pathways will be affected, initially resulting in a more positive threshold for action potential firing, hence hypo-excitability. Upon loss of CAMK2 in more subcellular locations, other pathways get involved, resulting eventually in a decrease in action potential firing, hence hyper-excitability. Interestingly, when applying 2 different CAMK2 inhibitors we did not find any change in excitability, indicating that acute inhibition in slices does not directly affect neuronal excitability. Thus, whether the shift from hypo- to hyper-excitability seen in our *Camk2a^f/f^*;*Camk2b^f/f^;Cag-Cre^ER^* mice is caused by a structural or kinetic function of CAMK2 or maybe a combination of both is subject for future research.

We observed a decrease in the frequency but not in the amplitude of miniature excitatory and inhibitory postsynaptic currents only 21 days after deletion of CAMK2A and CAMK2B, which is not seen in *Camk2a^−/−^* and *Camk2b^−/−^* mice. However, at 10 days after gene deletion, no changes in mPSCs are observed. This indicates that the changes in excitability are not caused by changes in inhibitory and excitatory input.

At 21 days, it is possible that the neuron tries to compensate for the synaptic loss of CAMK2A and CAMK2B and the accompanying decrease in mPSC input by decreasing the threshold for excitability. Loss of CAMK2B has already been associated with a decreased volume of spines and the conversion of mature spines into immature spines (Okamoto et al., 2007). Moreover, CAMK2 activity has been shown to be involved in unsilencing silent synapses (Liao et al., 2001). Our observation of more immature, filopodia-shaped synapses in the *Camk2a^f/f^*;*Camk2b^f/f^;Cag-Cre^ER^* mice indeed supports the idea that CAMK2 is important for synapse formation and/or maintenance and indicates that the neuron tries to compensate for the loss of CAMK2 at the synapse by increasing somatic excitability. However, we cannot exclude the opposite possibility, in which the neuron compensates for the hyperexcitability by decreasing mPSC frequency. Furthermore, it is still possible that the changes in mPSCs and in excitability are 2 independent mechanisms, caused by deletion of CAMK2A and CAMK2B simultaneously.

The mouse models used in our study have a different onset of gene deletion. The *Camk2a^f/f^;Camk2b^f/f^-CAG-Cre^ER^* mice develop normal CAMK2 expression up to the moment of Tamoxifen injections, usually starting at P21. However, *Camk2a^−/−^* and *Camk2b^−/−^* mice lack CAMK2A or CAMK2B from germline. This could mean that differences observed can also be explained by differences during (embryological) development. However, for CAMK2A it was recently shown that adult deletion results in similar LTP and hippocampal-dependent learning deficits as germline deletion (Achterberg et al., 2014). Furthermore taking into account the late onset of CAMK2A expression (P1) (Bayer et al., 1999), it is unlikely that CAMK2A plays a role during the embryological development. Its role in early postnatal development remains to be uncovered. CAMK2B starts to be expressed at E12.5 (Karls et al., 1992; Bayer et al., 1999), and is therefore more likely to play an important role early in development. Indeed, for locomotion, germline deletion of *Camk2b* is more disruptive than deletion of *Camk2b* in adulthood (Kool et al., 2016), indicating an important role for CAMK2B in development for locomotion If this is also true for other phenotypes remains to be investigated. However, it is still possible that the lack of effect we see in the *Camk2a^−/−^* and *Camk2b^−/−^* single mutants is not only due to compensation of the other CAMK2 isoform still being present, but through other compensatory mechanisms kicking in during early development, and that adult deletion of CAMK2A or CAMK2B in inducible single mutants would cause similar changes in excitability and unitary synaptic transmission as seen in the inducible double Camk2a/Camk2b mutants studied here.

Taken together our data show critical involvement in neuronal excitability as well as synapse formation and shed new light on the full spectrum of CAMK2 function in neurons. Additionally, our data indicates that acute deletion of a gene can uncover novel functions, which could remain unseen using conventional knockout methods.

## Acknowledgements

We thank Erika Goedknegt, Minetta Elgersma and Mehrnoush Aghadavoud Jolfaei for technical support and Gerard Borst and Diana Rotaru for comments on the manuscript. This research was supported by the Netherlands Organization for Scientific Research (NWO-Veni and Vidi project to G.v.W.).

## References

1. Achterberg KG, Buitendijk GHS, Kool MJ, Goorden SMI, Post L, Slump DE, Silva AJ, van Woerden GM, Kushner SA, Elgersma Y (2014) Temporal and Region-Specific Requirements of αCaMKII in Spatial and Contextual Learning. J Neurosci 34:11180– 11187.

2. Ashpole NM, Song W, Brustovetsky T, Engleman EA, Brustovetsky N, Cummins TR, Hudmon A (2012) Calcium/calmodulin-dependent protein kinase II (CaMKII) inhibition induces neurotoxicity via dysregulation of glutamate/calcium signaling and hyperexcitability. Journal of Biological Chemistry 287:8495–8506.

3. Barcomb K, Buard I, Coultrap SJ, Kulbe JR, O’Leary H, Benke TA, Bayer KU (2014) Autonomous CaMKII requires further stimulation by Ca2+/calmodulin for enhancing synaptic strength. FASEB J.

4. Bayer KU, Löhler J, Schulman H, Harbers K (1999) Developmental expression of the CaM kinase II isoforms: ubiquitous gamma- and delta-CaM kinase II are the early isoforms and most abundant in the developing nervous system. Brain Res Mol Brain Res 70:147–154.

5. Benson DL, Gall CM, Isackson PJ (1992) Dendritic localization of type II calcium calmodulin-dependent protein kinase mRNA in normal and reinnervated rat hippocampus. NSC 46:851–857.

6. Borgesius NZ, van Woerden GM, Buitendijk GHS, Keijzer N, Jaarsma D, Hoogenraad CC, Elgersma Y (2011) βCaMKII plays a nonenzymatic role in hippocampal synaptic plasticity and learning by targeting αCaMKII to synapses. J Neurosci 31:10141– 10148.

7. Burgin KE, Waxham MN, Rickling S, Westgate SA, Mobley WC, Kelly PT (1990) In situ hybridization histochemistry of Ca2+/calmodulin-dependent protein kinase in developing rat brain. J Neurosci 10:1788–1798.

8. Elgersma Y, Fedorov NB, Ikonen S, Choi ES, Elgersma M, Carvalho OM, Giese KP, Silva AJ (2002) Inhibitory autophosphorylation of CaMKII controls PSD association, plasticity, and learning. Neuron 36:493–505.

9. Erondu NE, Kennedy MB (1985) Regional distribution of type II Ca2+/calmodulin-dependent protein kinase in rat brain. J Neurosci 5:3270–3277.

10. Ghosh S, Reuveni I, Lamprecht R, Barkai E (2015) Persistent CaMKII Activation Mediates Learning-Induced Long-Lasting Enhancement of Synaptic Inhibition. J Neurosci 35:128–139.

11. Giese KP, Fedorov NB, Filipkowski RK, Silva AJ (1998) Autophosphorylation at Thr286 of the alpha calcium-calmodulin kinase II in LTP and learning. Science 279:870–873.

12. Hojjati MR, van Woerden GM, Tyler WJ, Giese KP, Silva AJ, Pozzo-Miller L, Elgersma Y (2007) Kinase activity is not required for alphaCaMKII-dependent presynaptic plasticity at CA3-CA1 synapses. Nat Neurosci 10:1125–1127.

13. Hund TJ, Koval OM, Li J, Wright PJ, Qian L, Snyder JS, Gudmundsson H, Kline CF, Davidson NP, Cardona N, Rasband MN, Anderson ME, Mohler PJ (2010) A β(IV)- spectrin/CaMKII signaling complex is essential for membrane excitability in mice. J Clin Invest 120:3508–3519.

14. Incontro S, Díaz-Alonso J, Iafrati J, Vieira M, Asensio CS, Sohal VS, Roche KW, Bender KJ, Nicoll RA (2018) The CaMKII/NMDA receptor complex controls hippocampal synaptic transmission by kinase- dependent and independent mechanisms. Nat Commun:1–21.

15. Isaacson JS, Scanziani M (2011) How Inhibition Shapes Cortical Activity. Neuron 72:231–243.

16. Karls U, Müller U, Gilbert DJ, Copeland NG, Jenkins NA, Harbers K (1992) Structure, expression, and chromosome location of the gene for the beta subunit of brain-specific Ca2+/calmodulin-dependent protein kinase II identified by transgene integration in an embryonic lethal mouse mutant. Mol Cell Biol 12:3644–3652.

17. Kim K, Lakhanpal G, Lu HE, Khan M, Suzuki A, Kato-Hayashi M, Narayanan R, Luyben TT, Matsuda T, Nagai T, Blanpied TA, Hayashi Y, Okamoto K (2015) A Temporary Gating of Actin Remodeling during Synaptic Plasticity Consists of the Interplay between the Kinase and Structural Functions of CaMKII. Neuron 87:813–826.

18. Klug JR, Mathur BN, Kash TL, Wang H-D, Matthews RT, Robison AJ, Anderson ME, Deutch AY, Lovinger DM, Colbran RJ, Winder DG (2012) Genetic inhibition of CaMKII in dorsal striatal medium spiny neurons reduces functional excitatory synapses and enhances intrinsic excitability. PLoS ONE 7:e45323.

19. Kool MJ, Onori MP, Borgesius NZ, van de Bree JE, Elgersma-Hooisma M, Nio E, Bezstarosti K, Buitendijk GHS, Aghadavoud Jolfaei M, Demmers JAA, Elgersma Y, van Woerden GM (2019) CAMK2-dependent signaling in neurons is essential for survival. Journal of Neuroscience:1341–18.

20. Kool MJ, van de Bree JE, Bodde HE, Elgersma Y, van Woerden GM (2016) The molecular, temporal and region-specific requirements of the beta isoform of Calcium/Calmodulin-dependent protein kinase type 2 (CAMK2B) in mouse locomotion. Sci Rep 6:26989.

21. Lee S-JR, Escobedo-Lozoya Y, Szatmari EM, Yasuda R (2009) Activation of CaMKII in single dendritic spines during long-term potentiation. Nature 458:299–304.

22. Liao D, Scannevin RH, Huganir R (2001) Activation of silent synapses by rapid activity-dependent synaptic recruitment of AMPA receptors. J Neurosci 21:6008–6017.

23. Lledo PM, Hjelmstad GO, Mukherji S, Soderling TR, Malenka RC, Nicoll RA (1995) Calcium/calmodulin-dependent kinase II and long-term potentiation enhance synaptic transmission by the same mechanism. Proc Natl Acad Sci USA 92:11175– 11179.

24. Mayford M, Bach ME, Huang YY, Wang L, Hawkins RD, Kandel ER (1996) Control of memory formation through regulated expression of a CaMKII transgene. Science 274:1678–1683.

25. Nelson AB, Gittis AH, Lac Du S (2005) Decreases in CaMKII activity trigger persistent potentiation of intrinsic excitability in spontaneously firing vestibular nucleus neurons. Neuron 46:623–631.

26. Ninan I, Arancio O (2004) Presynaptic CaMKII is necessary for synaptic plasticity in cultured hippocampal neurons. Neuron 42:129–141.

27. Okamoto KI, Narayanan R, Lee SH, Murata K, Hayashi Y (2007) The role of CaMKII as an F-actin-bundling protein crucial for maintenance of dendritic spine structure. Proceedings of the National Academy of Sciences 104:6418–6423.

28. Sametsky EA, Disterhoft JF, Ohno M (2009) Autophosphorylation of αCaMKII downregulates excitability of CA1 pyramidal neurons following synaptic stimulation. Neurobiology of Learning and Memory 92:120–123.

29. Silva AJ, Paylor R, Wehner JM, Tonegawa S (1992a) Impaired spatial learning in alpha- calcium-calmodulin kinase II mutant mice. Science 257:206–211.

30. Silva AJ, Stevens CF, Tonegawa S, Wang Y (1992b) Deficient hippocampal long-term potentiation in alpha-calcium-calmodulin kinase II mutant mice. Science 257:201– 206.

31. Thiagarajan TC, Piedras-Renteria ES, Tsien RW (2002) alpha- and betaCaMKII. Inverse regulation by neuronal activity and opposing effects on synaptic strength. Neuron 36:1103–1114.

32. Tobimatsu T, Fujisawa H (1989) Tissue-specific expression of four types of rat calmodulin-dependent protein kinase II mRNAs. J Biol Chem 264:17907–17912.

33. van Woerden GM, Hoebeek FE, Gao Z, Nagaraja RY, Hoogenraad CC, Kushner SA, Hansel C, De Zeeuw CI, Elgersma Y (2009) betaCaMKII controls the direction of plasticity at parallel fiber-Purkinje cell synapses. Nat Neurosci 12:823–825.

34. Varga AW, Yuan L-L, Anderson AE, Schrader LA, Wu G-Y, Gatchel JR, Johnston D, Sweatt JD (2004) Calcium-calmodulin-dependent kinase II modulates Kv4.2 channel expression and upregulates neuronal A-type potassium currents. Journal of Neuroscience 24:3643–3654.

35. Wang Z, Wilson GF, Griffith LC (2002) Calcium/calmodulin-dependent protein kinase II phosphorylates and regulates the Drosophila eag potassium channel. J Biol Chem 277:24022–24029.

36. Wei J, Zhang M, Zhu Y, Wang JH (2004) Ca2+–calmodulin signalling pathway up- regulates GABA synaptic transmission through cytoskeleton-mediated mechanisms. Neuroscience 127:637–647.

37. Zhou S, Yu Y (2018) Synaptic E-I Balance Underlies Efficient Neural Coding. Front Neurosci 12:46.

